# VitisC: Visualizing interested target via integrated scaffold of cytoophidium

**DOI:** 10.1101/2025.01.17.633574

**Authors:** Boqi Yin, Huan-Huan Hu, Jiale Zhong, Ji-Long Liu

## Abstract

Cryo-electron microscopy (cryo-EM) single-particle analysis has become a widely used technique for high-resolution structural determination of biological macromolecules and complexes. However, the determination of the structure of small molecule proteins remains a limitation of this technology. To address this issue, here we develop a novel approach termed Visualizing interested target via integrated scaffold of cytoophidium (VitisC). We use a filamentous structure formed by *Escherichia coli* CTP synthase (CTPS) as the scaffold, termed the scaffold of cytoophidium. Through artificial design and modification, small proteins can be attached to this symmetrical scaffold, which is very suitable for cryo-EM imaging. The formation conditions for stable filament structures of the fusion protein are optimized in vitro, and the three-dimensional structure of the fusion protein is reconstructed using cryo-EM single-particle analysis, achieving an overall resolution of 3.39 Å. Therefore, cryo-EM is successfully applied to visualize small proteins. VitisC not only demonstrates the feasibility of employing cytoophidia as scaffolds, but also provides an approach for high-resolution structural analysis of small proteins using Cryo-EM.

## INTRODUCTION

Over the past decade, cryo-electron microscopy (cryo-EM) single-particle analysis has advanced rapidly, with numerous technical breakthroughs. Cryo-EM has become a widely used technology for high-resolution structural analysis of biological macromolecular complexes. However, despite these advancements, the technology faces some challenges^1^. Achieving high resolution for proteins or complexes with small molecular weights using Cryo-EM single-particle analysis remains difficult. According to statistics from the Electron Microscopy Data Bank, structures of samples weighing less than 50 kDa account for only 1.1% of all published structures. This limitation arises from the inherently low contrast and poor signal-to-noise ratio of small molecular objects in frozen samples^2^. Furthermore, small molecular objects often lack distinct and recognizable structural features, which hinders initial image alignment and subsequent analysis^3^.

Statistics indicate that vast biologically important proteins fall outside the range of high-resolution structural determination achievable by this technique^4^. Overcoming the size limitations of Cryo-EM single-particle analysis is crucial. Currently, several research strategies have been proposed for small molecular proteins. One approach focuses on improving electron microscopy hardware and sample preparation techniques^5,6^. Another strategy involves attaching small molecular proteins to high molecular weight proteins or complexes, referred to as scaffolds^7,8^. This effectively integrates small molecular proteins into larger assemblies, enabling their structural elucidation as part of the whole complex. In recent years, innovative imaging scaffolds for cryo-EM have been developed, enabling the structural analysis of small proteins using this technique^9,10^. However, the rigidity of small proteins and the rapid modular application are still the two major challenges in this field^11^.

In this study, we develop an approach termed Visualizing interested target via integrated scaffold of cytoophidium (VitisC). We design a novel filamentous imaging scaffold, termed the scaffold of cytoophidia. The cytoophidium, first discovered in 2010 and formally named due to its serpentine shape ^12–14^, is a large assembly of metabolic enzymes formed within cells under environmental and metabolic regulation. In vitro, the filament structure formed by metabolic enzymes exhibits high symmetry and periodicity, enabling excellent 3D reconstruction from a limited number of particles. This structure also significantly alleviates the common issue of insufficient particle orientation, making it highly suitable for cryo-EM imaging.

By attaching mEGFP as a small protein to our engineered *Escherichia coli* CTPS scaffold, we successfully identify the conditions required for the fusion protein to form filaments in vitro. Using cryo-EM single-particle analysis, we achieve a filament structure with an overall resolution of 3.39 Å and successfully visualize the density of the small protein. VitisC demonstrates the feasibility of the scaffold of cytoophidium and present a novel strategy for the rapid application of imaging scaffolds. Furthermore, VitisC provides new insights into the potential applications of cytoophidia.

## RESULTS

### Designing the scaffold of cytoophidium

An optimal scaffold of cytoophidium must fulfill three key criteria. To begin with, the fusion protein should form a filament structure, ensuring that the attachment of the small protein does not disrupt the assembly of the filaments. In addition, the connection between the small protein and the scaffold of cytoophidium should be sufficiently rigid. Lastly, the scaffold should be easily modularized for versatility.

Building on prior studies of CTP synthase filaments, we observed that introducing a glutamate mutation after the N-terminal methionine allows stable filament formation in vitro under magnesium ion conditions. In this structure, electrostatic interactions between the N-terminal glutamate and the C-terminal His-tag play an important role in stabilizing the filament assembly (Fig. 1A). We infer that this interaction may help stabilize the otherwise flexible N-terminal region, which is beneficial to the rigid connection between the scaffold of cytoophidium and the small protein. Based on this discovery, we used *E. coli* CTP synthase as a scaffold and designed a linker to attach the small protein mEGFP at the N-terminus, constructing a series of fusion proteins (Fig. 1B). When the positioning of the small protein does not interfere with the interaction between the N-terminal glutamate and the C-terminal His-tag, the entire fusion protein is capable of forming a filamentous structure (Fig. 1C). This design also allows for convenient replacement of small proteins, avoiding complex screening processes and enabling rapid modular applications.

**Figure 1.**
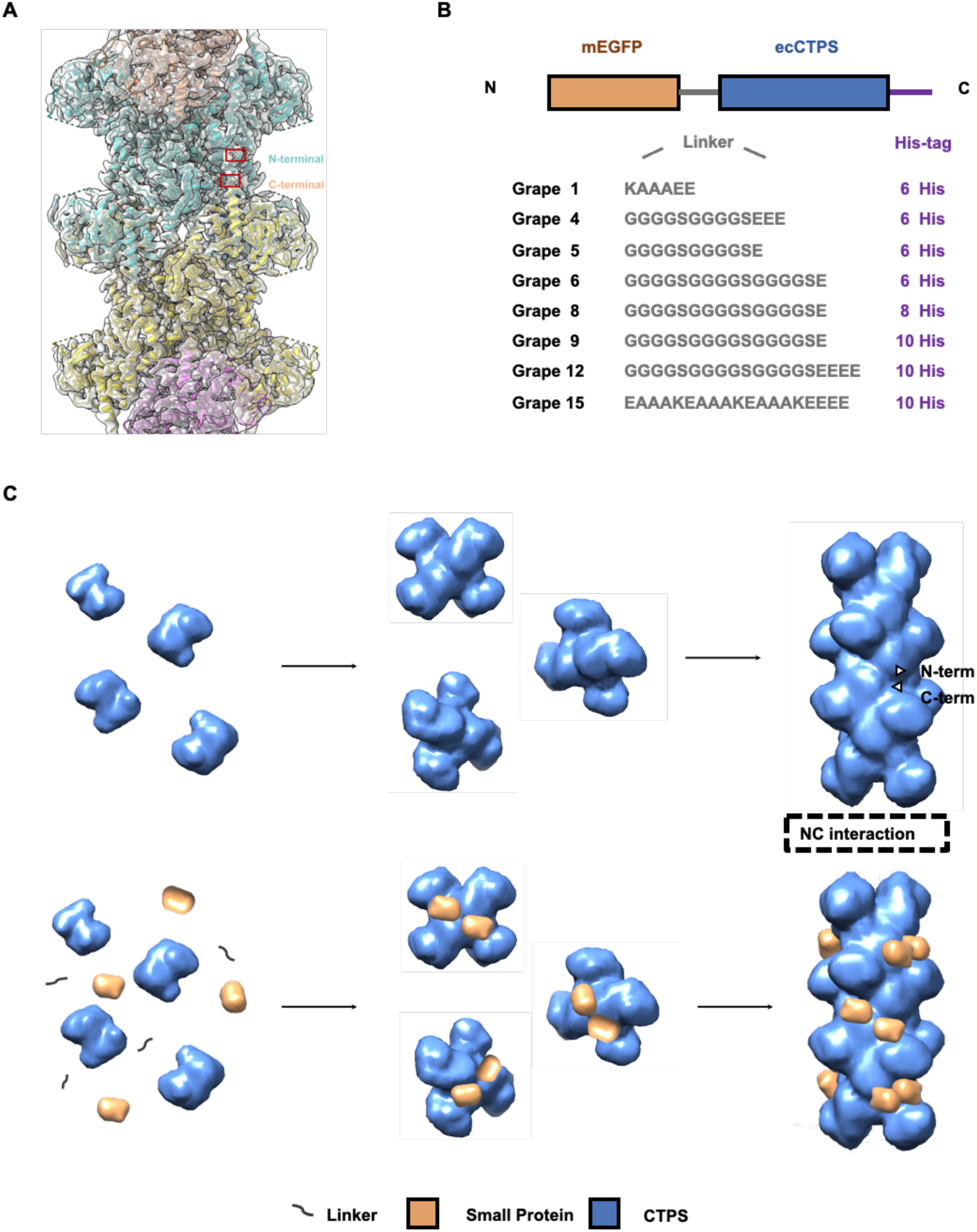
Designing the scaffold of cytoophidium and constructing fusion proteins. (A) Cryo-EM reconstruction of *E. coli* CTPS M13. The insertion of glutamic acid into the N-terminal enabled *E. coli* CTPS to form filaments under Mg²⁺ conditions, facilitated by potential electrostatic interactions between the N-terminal and C-terminal. (B) Design and construction of fusion proteins, referred to as the “Grape” series. (C) Model of the rigid connection between the scaffold of cytoophidium and the small protein. The small protein is stably connected to the N-terminal of CTP synthase without disrupting the interaction between the N-terminal and C-terminal of the tetramer. As a result, the fusion protein successfully forms a filamentous structure.

### Conditions for filament formation in vitro

Under APO, Mg²⁺, or product (Mg²⁺, CTP) conditions, the purified fusion protein was incubated to prepare negatively stained samples, and its morphology was observed using a 120 kV transmission electron microscope. Through continuous optimization of the linker and His-tag, it was found that the fusion proteins Grape9, Grape12, and Grape15 could form filaments under product conditions, but no filaments were observed under Mg²⁺ conditions (Fig. 2). Among the filamentous structures, Grape15 exhibited a long and straight morphology, making it the most suitable for subsequent analytical work. The conditions for forming stable filaments in vitro are as follows: 25°C under product conditions, with incubation in 25 mM Tris-HCl buffer (pH 8.0) for 30 minutes. No filaments of Grape15 were observed under APO or Mg²⁺ conditions. We hypothesize that the formation of filamentous structures by the fusion protein is influenced by the length of the linker and His-tag. Initially, our design focused on progressively lengthening the linker. However, when no filamentous structures were observed, we extended the His-tag and the experimental results demonstrated that filaments began to appear when the His-tag contained 10 histidine residues, indicating that the His-tag plays a crucial role in the formation of the filamentous structure in the fusion protein.

**Figure 2.**
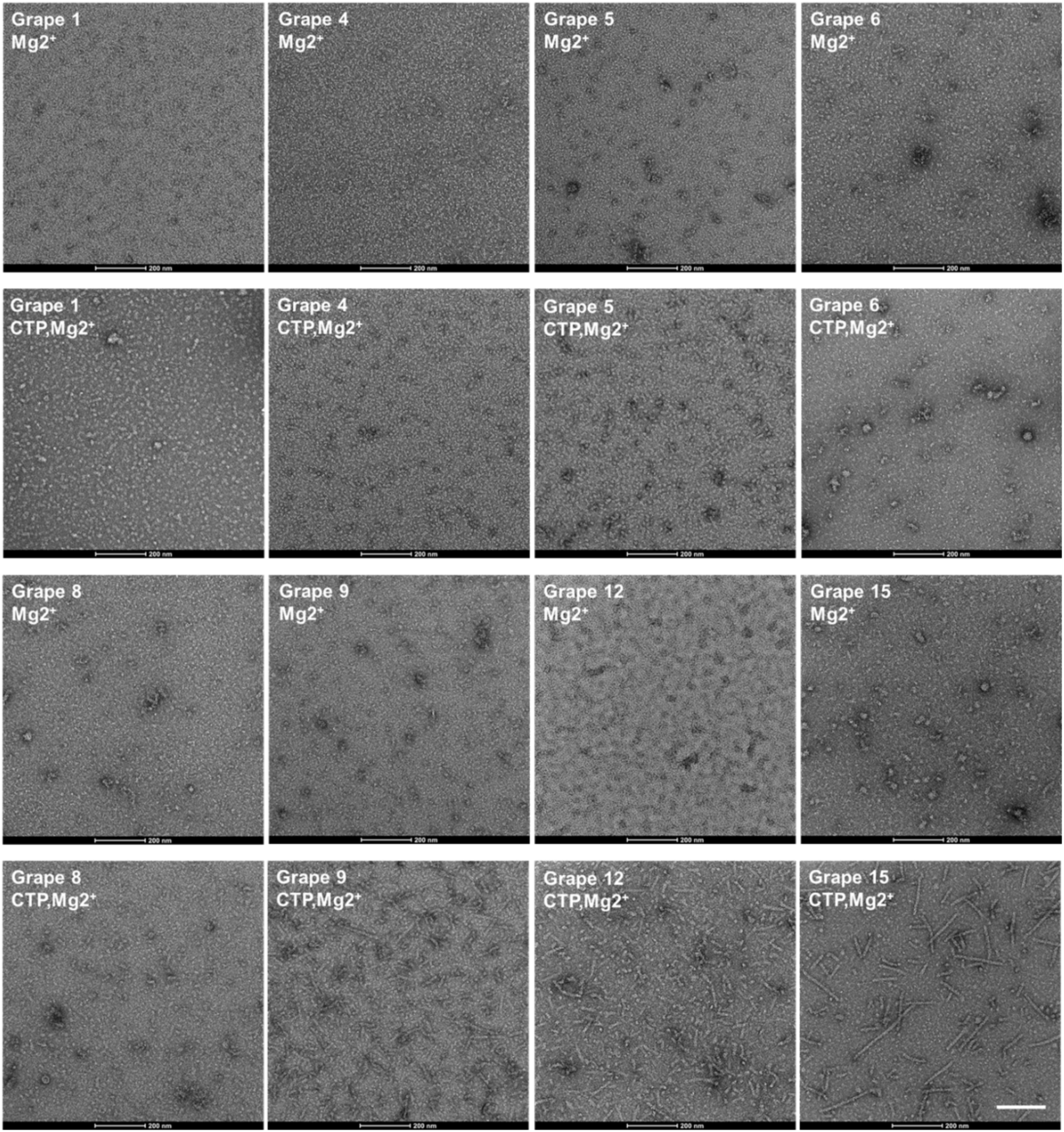
Representative negative staining results of fusion protein under Mg^2+^ condition and product condition in vitro. Under product conditions, fusion proteins Grape9, Grape12, and Grape15 were capable of forming filaments. However, no filament formation was observed under Mg²⁺ conditions. Protein concentration: 1 μM, Scale bar: 200 nm.

### Optimizing the connection between scaffold and small protein

Following the successful formation of stable filaments, Grape15 was selected for frozen sample preparation and structural analysis. The results revealed the absence of small protein density in the 2D classification (Fig. 3A), indicating that the connection between the small protein and the scaffold of cytoophidium required optimization. To achieve a more rigid model, the linker connecting the small protein to the scaffold was shortened, and a flexible sequence at the C-terminus of the mEGFP protein was truncated.

**Figure 3.**
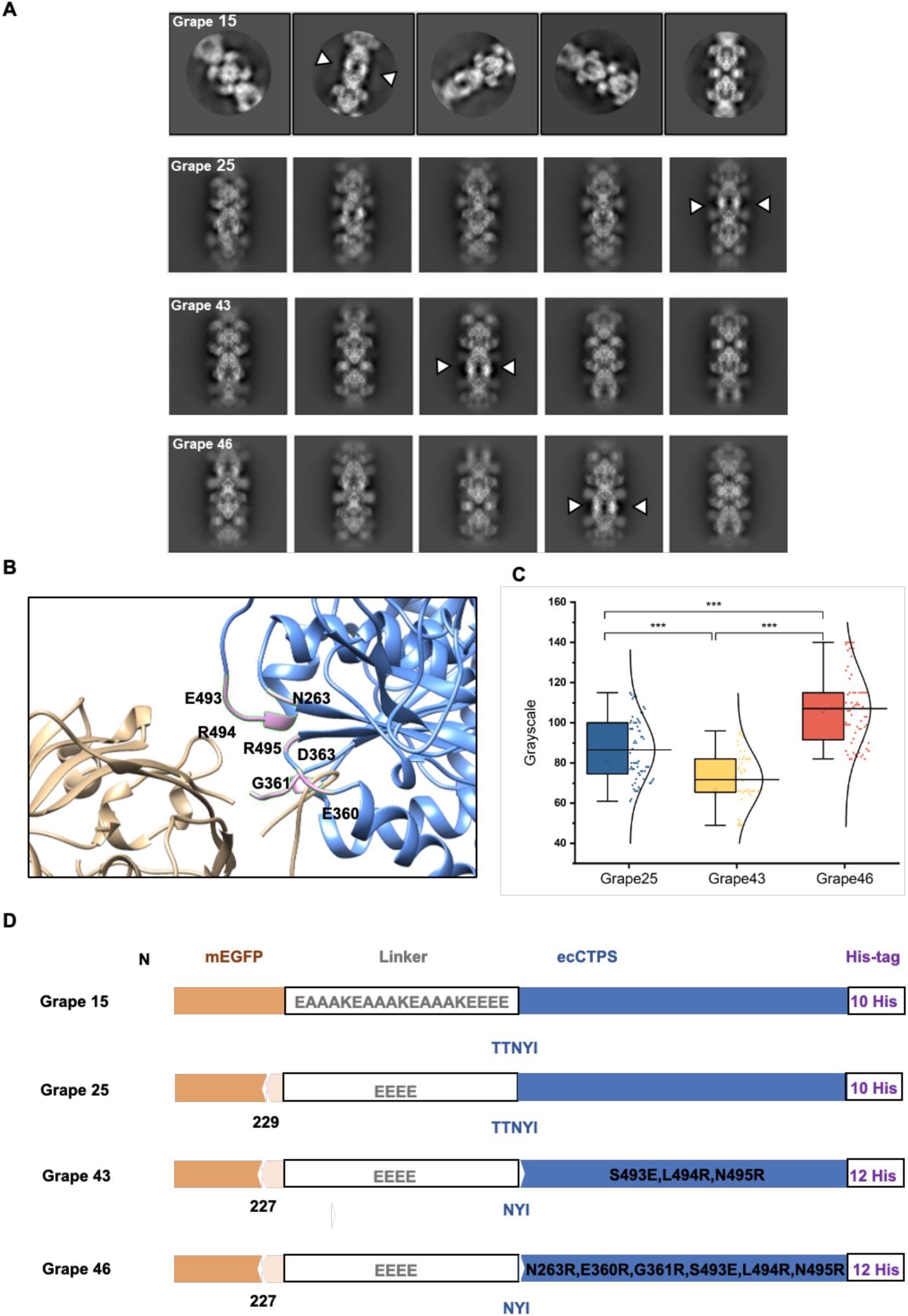
Optimization of the junction between the small protein and the scaffold of cytoophidium. (A) 2D classification results for Grape15, Grape25, Grape43, and Grape46. While Grape15 showed no visible small protein density, Grape25, Grape43, and Grape46 exhibited clear densities corresponding to the small proteins. The box sizes for the classifications are 380 pixels, 384 pixels, 360 pixels, and 360 pixels, respectively. (B) The analysis of the interaction interface between the scaffold and the small protein revealed specific amino acids on the scaffold that are likely to interact with the small protein. (C) Quantitative analysis of the density clarity of small proteins in representative 2D classifications of fusion proteins Grape25, Grape43, and Grape46 indicates that the small protein density in Grape46 is the most distinct, exhibiting the highest clarity and rigidity. (D) Protein sequence alignment of Grape15, Grape25, Grape43, and Grape46.

Based on an analysis of the existing structural data, specific amino acids on the scaffold that may interact with the small proteins were mutated (Fig. 3B). We aim to design amino acids “grips” on the scaffold to enhance the stabilization of small proteins. Furthermore, to preserve the integrity of the filament structure, the length of the C-terminal His-tag was extended. Subsequent steps included purification, negative staining (Fig. S1A), preparation of frozen samples, and data collection of the optimized fusion protein.

Compared to Grape15, the optimized variants—Grape25, Grape43, and Grape46 exhibited clear localization and density of small proteins in the 2D classification results (Fig. 3A). By using the grayscale analysis function of ImageJ software, the clarity of small protein density in 2D classification results was quantitatively assessed (Fig. S1B). A higher grayscale value indicates greater contrast between the small protein and the background noise in the cryo-sample, reflecting increased rigidity in the connection between the small protein and the scaffold of cytoophidium. The results demonstrate that the small protein density of Grape46 is the clearest (Fig. 3C), and the best 3D reconstruction results can be obtained, which is also confirmed by subsequent data processing.

This method offers a valuable approach for efficiently screening the scaffold of cytoophidium in future applications, significantly reducing the time required for data collection and 3D reconstruction. A comparison of the protein sequences is provided (Fig. 3D).

### Cryo-EM analysis of Grape46 filaments

Here we present the results of the freezing data analysis for Grape46(Fig. S2). In the observed frozen samples, Grape46 proteins formed highly regular filaments (Fig. 4A). A filamentous map comprising three tetramers was reconstructed, achieving an overall resolution of 3.39 Å (Fig. 4B). Local resolution estimation of the 3D electron density map for the Grape46 fusion protein revealed that the resolution of the scaffold of cytoophidium reached approximately 3 Å, while the surrounding small protein densities ranged from 4 to 6.5 Å (Fig. 4C). Benefiting from the scaffold of cytoophidium, the collected fusion protein data exhibited more complete angular information (Fig. 4D).

**Figure 4.**
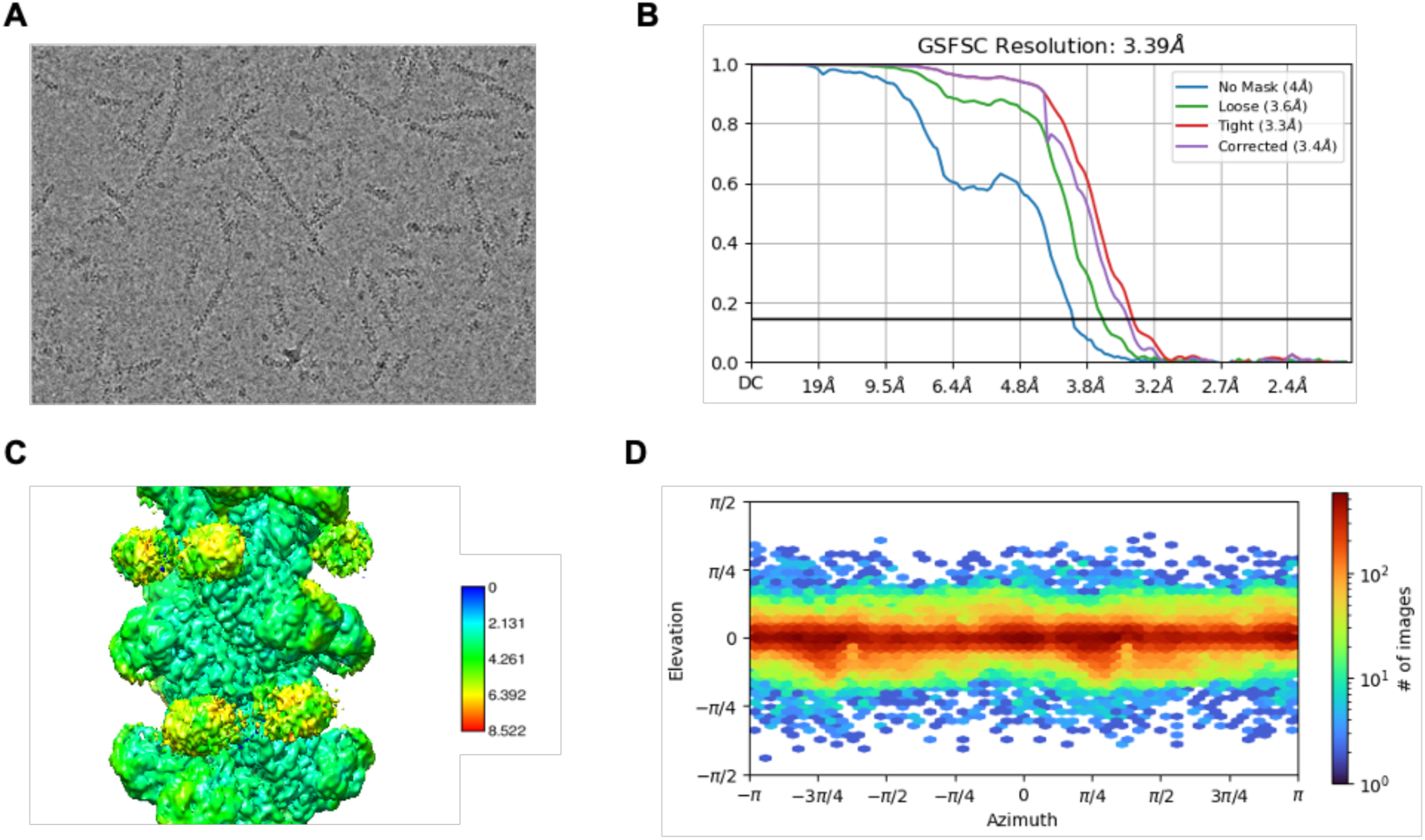
Representative results from 300 kV cryo-EM data processing of Grape46. (A) Representative 300 kV cryo-EM image of Grape46 after motion correction. (B) Gold-standard Fourier shell correlation (FSC) curves for Grape46 after 3D refinement. (C) Local resolution estimation of the 3D density map of the Grape46 fusion protein. (D) Angular distribution of particle orientations in the refined Grape46 reconstruction.

In the 3D electron density map of Grape46, the His-tag located at the C-terminal of the fusion protein was tightly connected to its adjacent tetramer, while the NC-terminal junction demonstrated a well-defined small protein density (Fig. 5A). The electron density map further emphasized the critical role of the C-terminal His-tag in fibrous structure formation (Fig. S3). Under product conditions, CTPS tetramers relied primarily on the His-tag to assemble into filament structures. When only eight histidine residues were present, the C-terminal of one tetramer was too distant from the adjacent tetramer to interact, consistent with our earlier experiments screening conditions for filaments formation.

**Figure 5.**
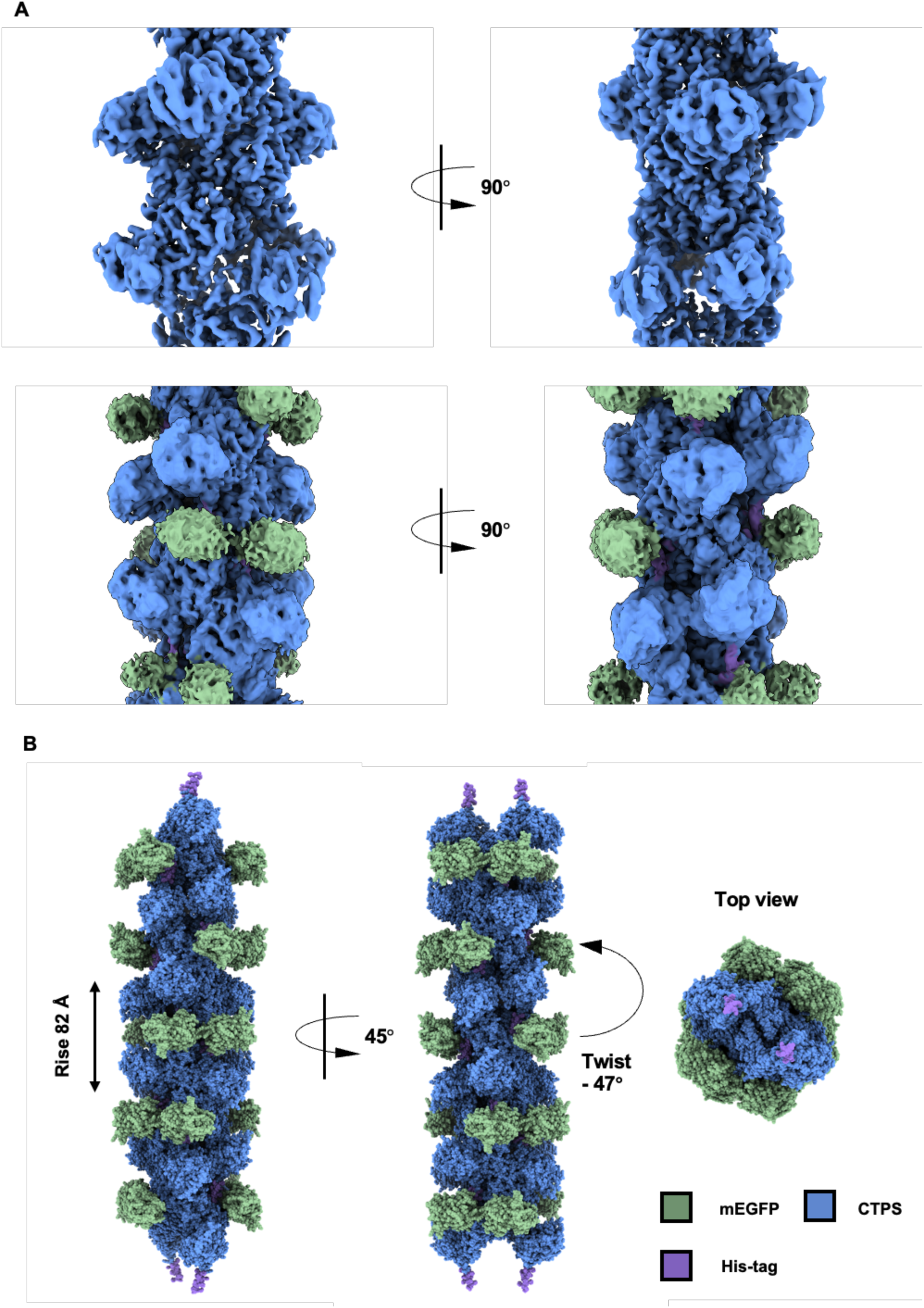
Overall 3D structure of Grape46 filaments. (A) The density map of the Grape46 fusion protein is displayed with the CTPS scaffold and small protein mEGFP highlighted in distinct colors. (B) A proposed spherical model of Grape46 filaments is presented from three different perspectives.

An atomic model of the Grape46 fusion protein was constructed using Coot, resulting in a filament structure consisting of five stacked tetramers. The stacking of tetramers into a single fiber was mediated by interactions between the C-terminal His-tag and the adjacent tetramer. The rise of the entire fusion protein fiber is approximately 82 Å, and the twist is about −47° (Fig. 5B).

### Interface between small protein and the scaffold of cytoophidium

When the fusion protein tetramer model was fitted into the electron density map, the overall density aligned well with the model (Fig. 6A). The effectiveness of the amino acids “grips” designed on the Grape46 scaffold was analyzed using ChimeraX visualization. Specifically, E493 formed electrostatic interactions with histidine on the small protein, R494 created a salt bridge with glutamic acid on the linker, and R495 formed a salt bridge with aspartic acid on the small protein, effectively stabilizing the small protein (Fig. 6B).

**Figure 6.**
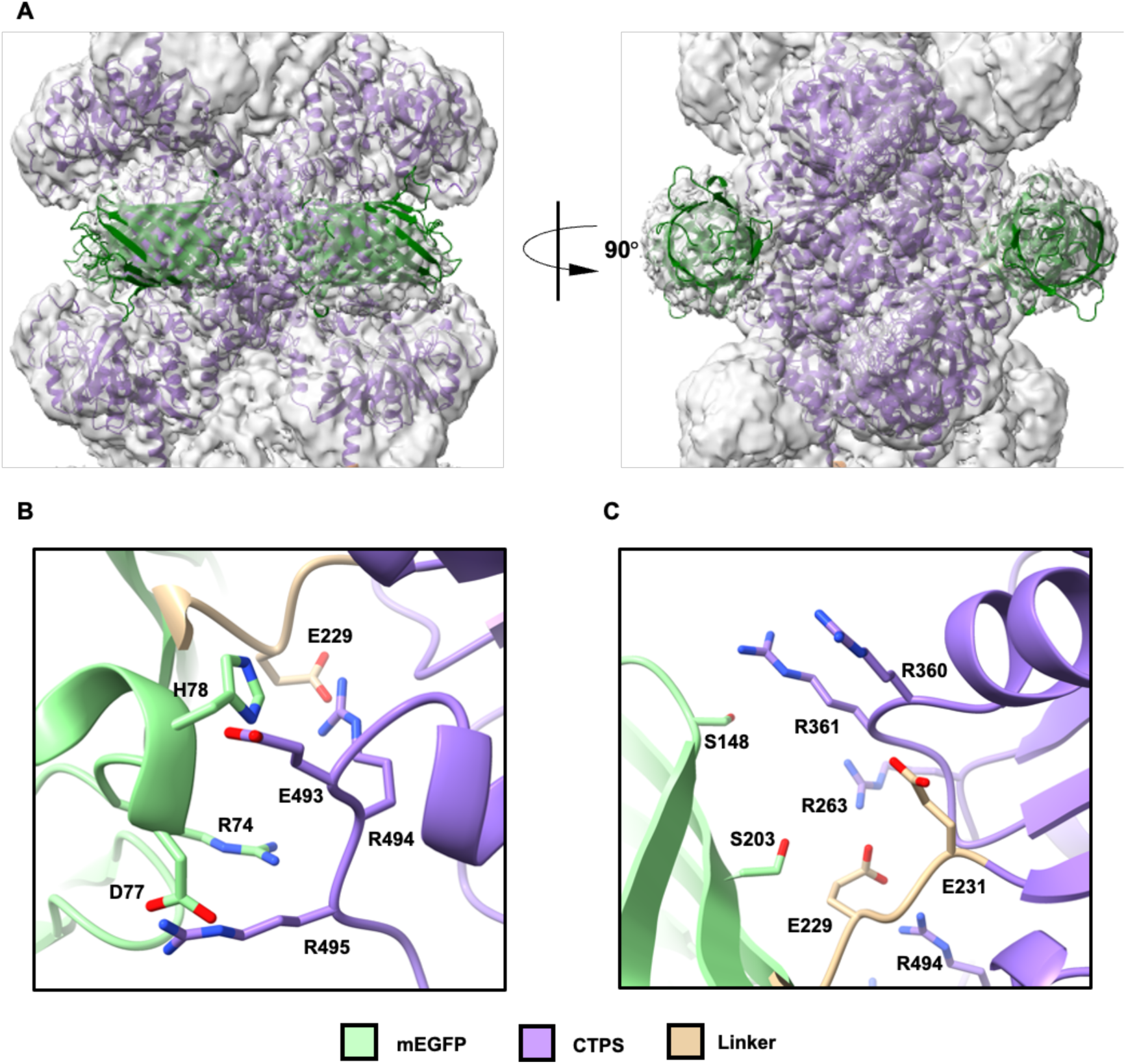
Cryo-EM Structure of Grape46. (A) A refined atomic model of the Grape46 tetramer fitted into the cryo-EM density map. (B) The interaction interface between the CTPS scaffold and mEGFP reveals that mutated amino acids on the CTPS and the linker contribute to stabilizing the small proteins.

Additionally, R361 established a hydrogen bond with E231 on the linker, while R263 engaged in electrostatic interactions with E229 on the linker, similar to R494, thereby stabilizing the connection between the N-terminal of the scaffold of cytoophidium and the small protein and reducing the flexibility of the small protein. R360, however, appeared to be an ineffective mutation (Fig. 6C).

Overall, the amino acids “grips” we designed serve a dual purpose: they stabilize the connection between the N-terminal of the scaffold and the small protein, thereby reducing the flexibility of the small protein relative to the scaffold, while also directly interacting with the small protein to enhance its rigidity significantly.

## DISCUSSION

We have successfully developed VitisC to visualize the 3D electron density of the small proteins using cryo-EM. We investigate the conditions under which the scaffold of cytoophidium can maintain filament structures after connecting with small proteins. The development of VitisC demonstrates that a novel filament imaging scaffold has been successfully designed and is ready for rapid application.

In addition, our study provides a clear direction for scaffold optimization, including the design of additional amino acid “grips”. Furthermore, we find that His-tags, traditionally used in protein purification, can also serve as critical interaction points for metabolic enzyme filamentation.

Notably, previously published data have identified more than 60 proteins capable of forming cytoophidia ^15–17^, many of which exhibit high symmetry and periodicity. For example, phosphoribosyl pyrophosphate synthase (PRPS) can form cytoophidia and metabolic filaments, and their high-resolution structures have been analyzed^18–20^.

Looking ahead, there is significant potential to update VitisC by exploring and designing additional scaffold of cytoophidium with promising applications. Beyond small protein structure analysis, these scaffolds can serve broader purposes in medical and industrial fields, such as protein proximity labeling, drug screening, and the development of nanomaterials.

## METHODS

### Construction of fusion proteins

The fusion proteins were constructed using *E. coli* CTP synthase (CTPS) as the scaffold and a mutant enhanced green fluorescent protein (mEGFP, A206K) as the small protein component. Both genes were cloned into a modified pRSFDuet vector with a C-terminal His-tag using a one-step cloning method (ClonExpress II One-Step Cloning Kit, Vazyme # C112). The scaffold, linker, and His-tag sequences were optimized through site-directed mutagenesis. DNA sequencing was performed to verify the accuracy of the constructed plasmid.

### Expression and purification of fusion proteins

The fusion proteins were expressed in *E. coli* Transetta (DE3) cells. The cells were initially cultured in auto-induction medium at 37°C with shaking at 220 rpm until the OD600 reached 0.5-0.8. The culture was then transferred to 16°C and continued for 16 hours at 150 rpm. Afterward, the cells were harvested by centrifugation at 4000 rpm for 10 minutes at 4°C. The collected cells were lysed by sonication in lysis buffer (50 mM Tris-HCl, pH 8.0, 500 mM NaCl, 10% glycerol, 20 mM imidazole, 1 mM PMSF, 5 mM β-mercaptoethanol, 2 μg/ml leupeptin, 2 μg/ml pepstatin, 1 mM benzamidine). The lysate was then centrifuged at 15,000 rpm for 50 minutes at 4°C. The supernatant was incubated with Ni-NTA agarose beads (Qiagen), pre-equilibrated with lysis buffer, for 1 hour at 4°C. The mixture was transferred to a column, washed with lysis buffer containing 40 mM imidazole, and the target protein was eluted with an elution buffer (50 mM Tris-HCl, pH 8.0, 500 mM NaCl, 250 mM imidazole, 5 mM β-mercaptoethanol). The eluted protein was further purified by gel filtration chromatography using a HiLoad 16/600 Superdex 200 pg column (GE Healthcare) or a Superose 6 Increase 10/300 GL column (Cytiva), both equilibrated with storage buffer (25 mM Tris-HCl, pH 8.0, 150 mM NaCl). The peak fractions were analyzed by SDS-PAGE after denaturation at 98°C. Finally, the target protein was concentrated, aliquoted, rapidly frozen in liquid nitrogen, and stored at −80°C.

### Negative staining electron microscopy

Under APO, Mg²⁺ (10 mM MgCl₂), or product conditions (10 mM MgCl₂, 2 mM CTP), the purified fusion protein was added to a protein activity buffer (25 mM MES, pH 6.0 for Grape43 and Grape46; 25 mM Tris-HCl, pH 8.0 for others) and incubated at 25°C for 30 minutes. The incubated fusion protein samples were then applied to glow-discharged copper grids with continuous carbon film, washed twice with deionized water, and stained with 1% uranium formate. The morphology and structure of the fusion protein were observed using a 120 kV transmission electron microscope (Talos L120C, ThermoFisher, USA).

### Cryo-EM grid preparation and data collection

The purified fusion protein was added to an activity buffer containing 10 mM MgCl₂ and 2 mM CTP (25 mM Tris-HCl, pH 8.0, for Grape15 and Grape25; 25 mM MES, pH 6.0, for Grape43 and Grape46). The final protein concentration was adjusted to 4–6 μM, and the sample was incubated at 25°C for 30 minutes prior to vitrification. The fusion protein sample was applied to an H₂/O₂ glow-discharged ANTcryo holey support film (Au300-R1.2/1.3), blotted for 3.5 seconds using a Vitrobot (Thermo Fisher) at 4°C and 100% humidity, and then rapidly plunged into liquid ethane cooled by liquid nitrogen. The sample was subsequently transferred to liquid nitrogen for storage.

The images were acquired using a 300 kV Titan Krios G3 (FEI) transmission electron microscope equipped with a K3 Summit direct electron detector (Gatan) operating in super-resolution mode. Automated data collection was performed using SerialEM^21^ at a magnification of 22,500×, resulting in a pixel size of 1.06 Å. The defocus range was set between 1.0 μm and 2.5 μm. For Grape15 and Grape25, the total dose was 60 e⁻/Å², subdivided into 50 frames at 2.8s exposure, while for Grape43 and Grape46, the total dose was 50 e⁻/Å², subdivided into 40 frames at 2.5 s exposure.

### Image processing

For Grape15, image processing was performed using RELION3.3^22,23^, which included drift correction with MotionCor2^24^ and contrast transfer function (CTF) estimation using CTFFIND4^25^. Images with an estimated resolution better than 4 Å were selected for particle picking. Initially, a subset of particles was manually selected, followed by several rounds of 2D classification to generate templates. These templates were subsequently used for automated particle picking across all images. After multiple rounds of 2D classification, the final results revealed distinct 2D classes, including two tetramers.

For Grape25, Grape43, and Grape46, image processing was carried out using CryoSPARC^26^. Initial preprocessing of the images involved patch motion correction and patch CTF estimation, and only images with an estimated resolution better than 3.8 Å were selected for further analysis. First, several dozen image stacks were manually selected, and particles were picked using 2D templates generated from the filamentous structure of wild-type *E. coli* CTPS (PDB: 5U3C)^27^. After several rounds of 2D classification, the resulting 2D classes were used by topaz train. The particles identified through Topaz train were then subjected to multiple rounds of 2D classification, ultimately yielding 2D classification results containing three tetramers.

For Grape46, an Ab-Initio Reconstruction was performed to generate an initial volume, which was subsequently refined using homogeneous reconstruction and non-uniform reconstruction. The final 3D map was further improved through CTF refinement to enhance resolution. Local resolution estimation was performed to determine the final resolution of the map. To further investigate the heterogeneity of the small proteins, we performed D2 symmetry expansion on the particles following CTF refinement. The box size was then reduced from 360 pixels to 180 pixels, concentrating on the central tetramer, and the particles were re-extracted. After multiple rounds of 2D classification, we performed consensus refinement with C1 symmetry for the particles. Next, a mask was created for a region containing a small protein and a CTPS monomer, and focused 3D classification was applied. From the six resulting classes, three were selected for subsequent 3D refinement.

### Model building

The original mutant *E. coli* CTPS monomer structure, along with the linker and small protein structures, were obtained from the AlphaFold2 database^28^. Using UCSF Chimera^29^, these structures were fitted into the final 3D reconstruction map. The tetramer structure of the fusion protein was subsequently constructed and refined in Coot^30^.The structural visualization was performed using UCSF ChimeraX^31^.

### Quantitative analysis of electron density in small protein structures

The 2D classification results of Grape25, Grape43, and Grape46 were sorted based on resolution, and the top 16 classes were selected for further analysis. These selected 2D images were imported into ImageJ and converted into 8-bit grayscale images. The XY coordinates were extracted, and the grayscale value data was imported into Origin software for analysis. The maximum grayscale value of each small protein region in each 2D class was calculated. The resulting values for Grape25, Grape43, and Grape46 were 62, 63, and 64, respectively. Statistical analysis of the data was performed using one-way ANOVA to determine significant differences between the groups.

## ACKNOWLEDGMENTS

EM data were collected at the ShanghaiTech Cryo-EM Imaging Facility. We thank the Molecular and Cell Biology Core Facility at the School of Life Science and Technology, ShanghaiTech University for providing technical support. This work was supported by grants to J.-L.L. from National Natural Science Foundation of China (number 32350710195) and UK Medical Research Council (grant numbers MC_UU_12021/3 and 474 MC_U137788471).

## SUPPLEMENTARY FIGURES S1-S3 AND FIGURE LEGENDS

**Figure S1.**
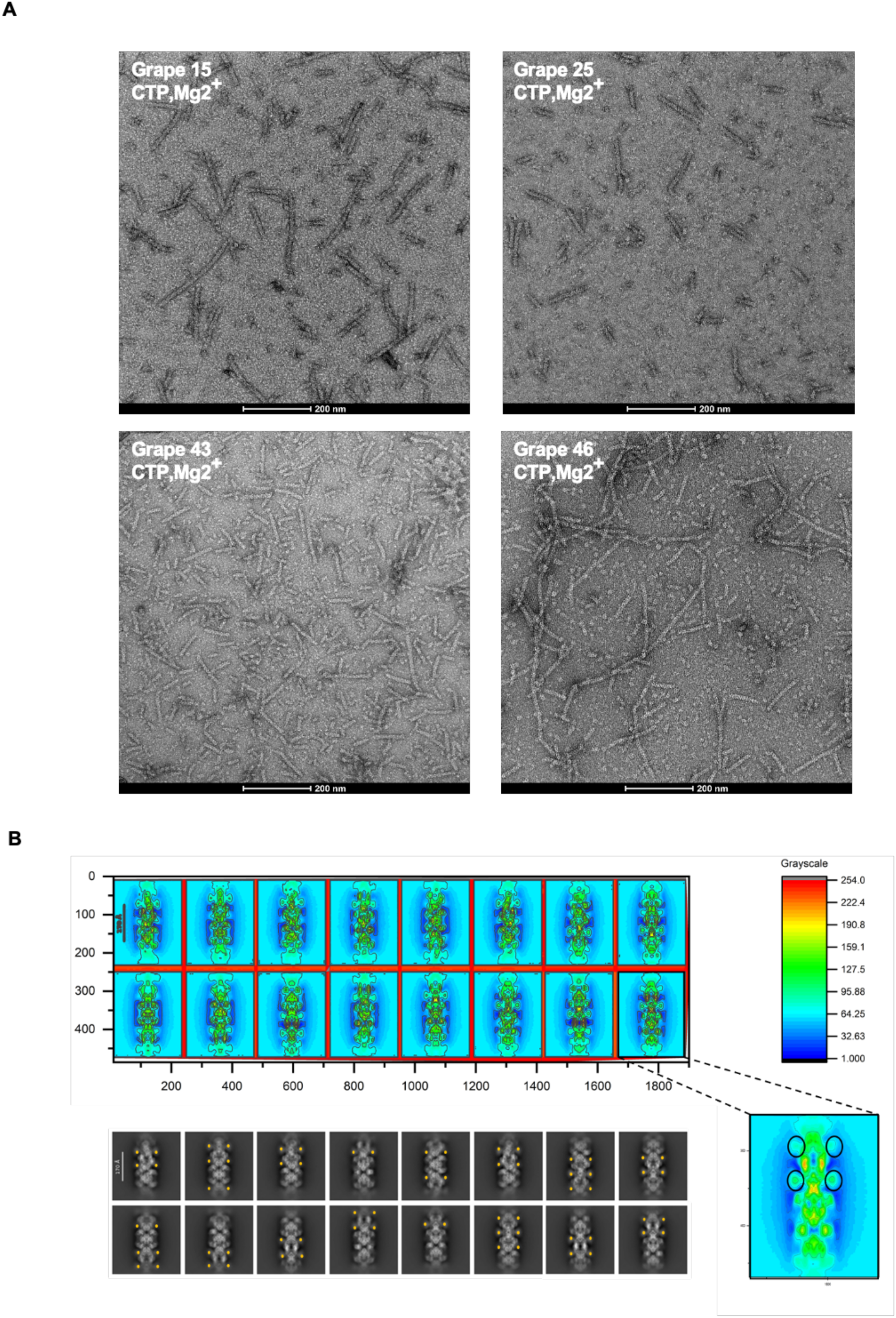
Negative staining and grSyscale analysis. (A) Negative staining results of fusion proteins Grape15, Grape25, Grape43, and Grape46 under product conditions in vitro. (B) The grayscale analysis of the small protein region is represented by Grape25. For each 2D classification result, the highest grayscale value within the small protein region was measured. A total of 16 types of 2D results were analyzed for Grape25, Grape43, and Grape46, yielding values of 68, 52, and 64, respectively.

**Figure S2.**
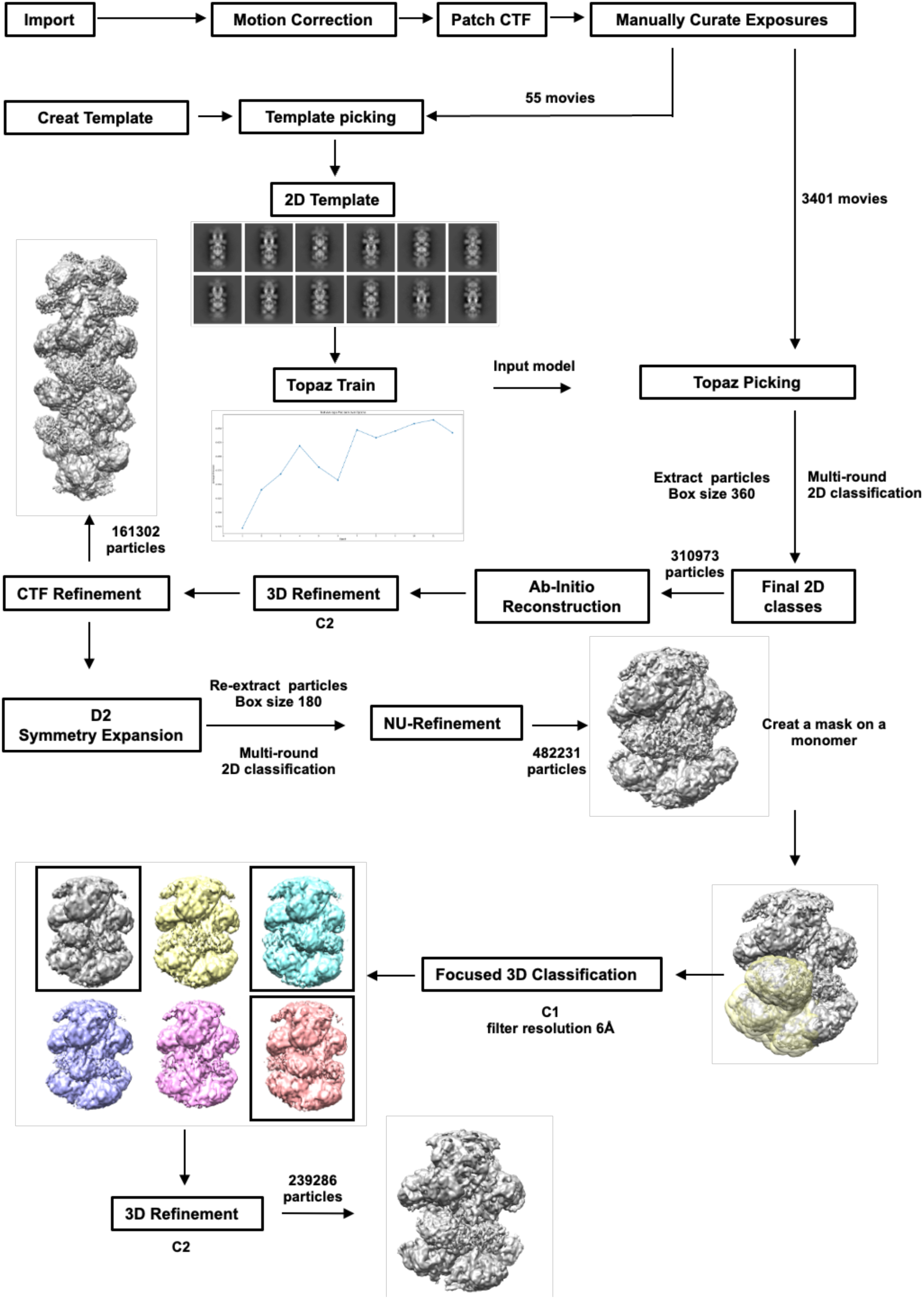
Data processing of Grape46 and focused refinement of small protein.

**Figure S3.**
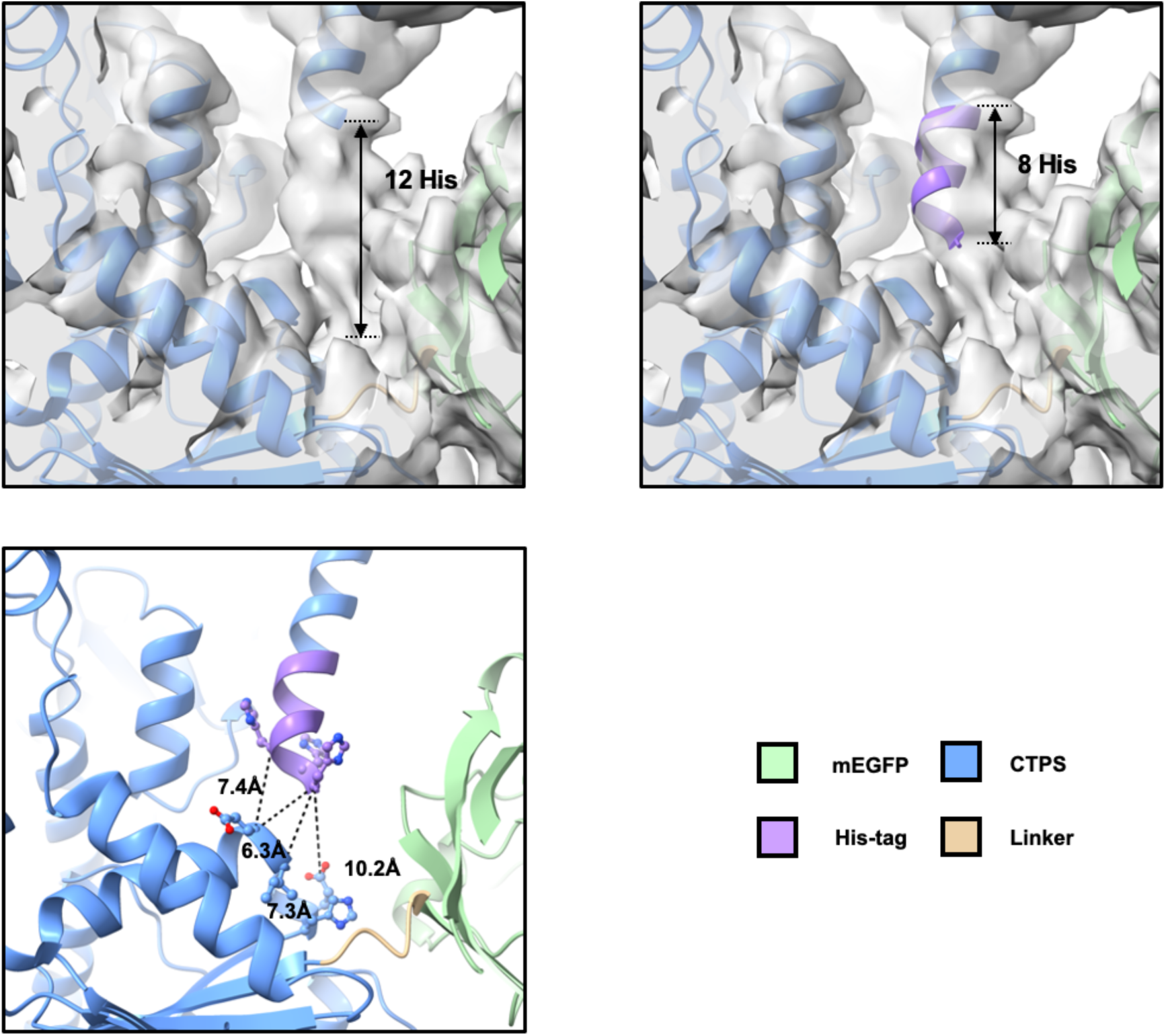
Analysis of the interaction interface of the tetramer in the filamentous structure of the Grape46 fusion protein.

